# Chitin degradation by *Synechococcus* WH7803

**DOI:** 10.1101/2023.06.07.544114

**Authors:** Giovanna Capovilla, Kurt G. Castro, Silvio Collani, Sean M. Kearney, David M. Kehoe, Sallie W. Chisholm

## Abstract

Chitin is an abundant, carbon-rich polymer in the marine environment. Chitinase activity has been detected in spent media of *Synechococcus* WH7803 cultures – yet it was unclear which specific enzymes were involved. Here we delivered a CRISPR tool into the cells via electroporation to generate loss-of-function mutants of putative candidates and identified ChiA as the enzyme required for the activity detected in the wild type.

## Main

The marine cyanobacterium *Synechococcus* is broadly distributed in the marine environment and is the most abundant group of phytoplankton globally in terms of total biomass^1^. As such, it contributes significantly to ocean primary productivity and the ocean carbon cycle. While these bacteria are considered primarily phototrophic and free-living, *Synechocococcus* strains possess chitin degradation genes and can switch from their canonical planktonic lifestyle to living in particles, including chitin^2^.

Chitin, an insoluble polymer of β1,4-linked N-acetylglucosamine (GlcNAc), is primarily derived from arthropod exoskeletons and serves as an important carbon and nitrogen source for marine microbial consortia^3–5^. To utilize this carbon source, bacteria must degrade chitin into soluble oligosaccharides via the action of enzymes defined as chitinases, which are divided into categories based on their activity^6^. Endochitinases are chitinases that cleave within the polymer strand of chitin, while exochitinases cleave terminal disaccharides from chitin oligosaccharides^6^. The latter are further characterized as chitobiosidases if they cleave dimeric units of GlcNAc from the non-reducing terminal of the polymer or β-N-acetylglucosaminidase if they cleave GlcNAc monomers^7^.

Both extracellular endochitinase and chitobiosidase activity were detected in cell-free supernatants of axenic *Synechococcus* WH7803 cultures^2^, indicating that the cells secrete active chitinases. However, the specific enzymes involved were unknown. Here we identify the genes responsible for chitin degradation and their role while broadening the toolbox available for *Synechoccoccus* genetic manipulation, demonstrating that electroporation is a reliable strategy to deliver gene editing tools.

We used bioinformatics to identify candidate chitinase genes. A putative chitinase gene in the *Synechococcus* WH7803 genome is WH7803_2068, which we refer to as *chiA* henceforth. ChiA contains a Beta-glycosidase of family GH18 listed as a possible chitinase and two N-terminal carbohydrate-binding modules of the CBM2^8^ family (Fig.S1a). We also identified two other proteins of interest – WH7803_2345 and WH7803_2069 – which contain two peripheral CBM2 modules and one central CBM2 module, respectively, with close homology to those in ChiA (Fig.S1a). Their relative position in the *Synechococcus* WH7803 genome is shown in Fig.1a and Fig.S2a. To determine whether the genes of interest respond to adding chitin to the media, we used qPCR to measure their expression in *Synechococcus* WH7803 cultures grown with and without chitosan or colloidal chitin. The three genes were expressed under all conditions, and their expression increased after adding chitin to the samples, but these increases were not statistically significant (Fig.S1b,c). Consistent with this observation, ChiA was abundant in a previous proteomic analysis even without chitin addition to the growth medium^8^. Similarly, expression of *chiA* genes is also constitutively expressed in *Prochlocococcus* MIT9303^2^ and other organisms such as diatoms^9^.

**Fig.1:**
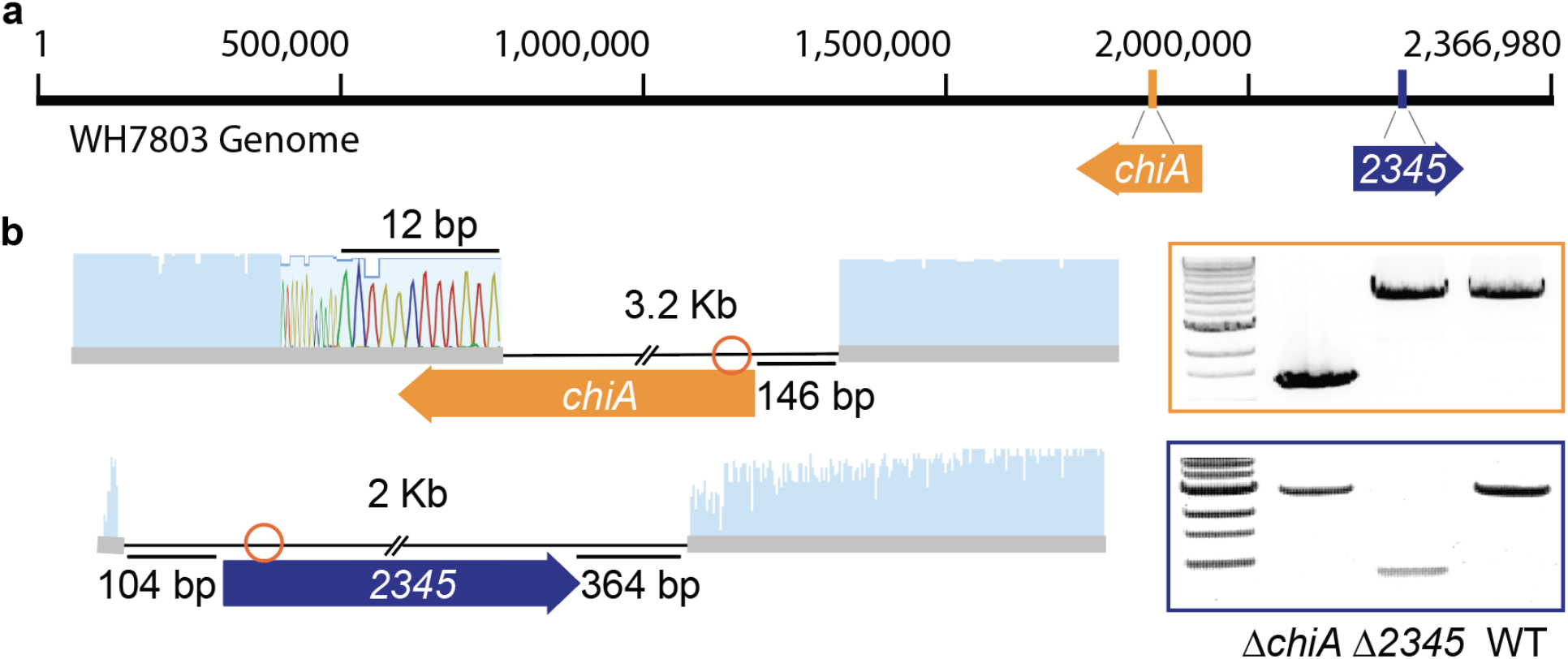
Mutants lacking *chiA* or *2345* obtained with a CRISPR-Cpf1 approach. **a**, Cartoon representation of the WH7803 genome and the relative positions of the genes of interest. **b**, Schematic representation of the edited cell lines obtained with the CRISPR-Cpf1 tool. Sanger sequences show the details of each deletion. Orange circles show the location of the PAM sites. PCR products indicate the length of each amplification using primers listed in Table S2, which are designed outside the homologous template regions.

To investigate the contribution of each of the three genes to chitin degradation, we designed an approach to obtain and test loss-of-function mutant lines. We employed a CRISPR-Cpf1 plasmid successfully used in freshwater cyanobacteria^10^ to make targeted deletions of each gene. This plasmid contains the CRISPR-Cpf1 cassette, a guide RNA to target a double-strand break on the gene of interest and a homologous repair template that cells can use to repair the DNA via homologous recombination. To deliver the engineered CRISPR plasmids (Table S1), we used an electroporation protocol^11^ with modifications (see methods) rather than a conjugation method, simplifying the recovery and purification of transformants. This work provides a new strategy for modifying cyanobacterial genomes when conjugation is unsuccessful or inefficient, as in the closely related species *Prochlorococcus*^*11*^.

The selection of fully edited lines was hampered by polyploidy in WH7803, which carries 3-4 genome copies^12–14^. Several rounds of plating and dilution-to-extinction with selection pressure were required to obtain fully segregating mutants. We ultimately obtained fully edited lines lacking *chiA* or *WH7803_2345*, which we call here *ΔchiA* and *Δ2345*, respectively (Fig.1b). Mutants were tested via qPCR to measure the level of expression of the targeted gene (Fig.2c-d), *i*.*e*., to determine if they were true knock-outs. We also obtained a deletion in *WH7803_2069* (Fig.S2), but we were unsuccessful in isolating a fully edited line, so we suspect that 2069 may be beneficial for growth in laboratory conditions. However, the mutant line obtained, *Δ2069*, shows a significantly lower expression of *2069* than the wild type (Fig.S2c), thus, we included it in our analysis, considering it a knock-down line.

**Fig.2:**
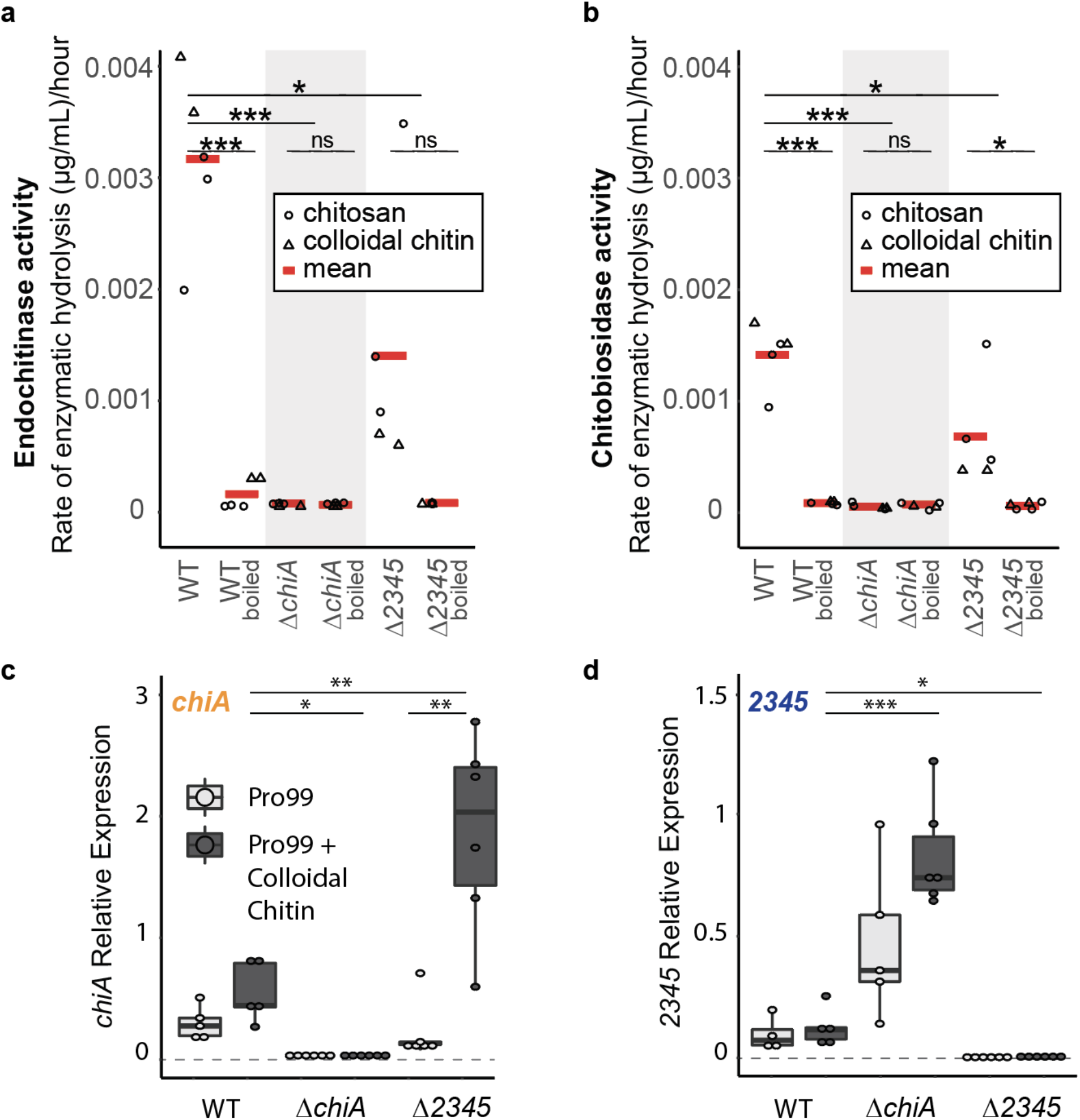
Rate of chitin degradation activity in mutants lacking ChiA or 2345 in comparison to wild-type WH7803. **a-b**, Endochitinase and Exochitinase (chitobiosidase) activities measured in wild-type and mutant lines spent media amended with either colloidal chitin or chitosan to a final concentration of 56 μg/ml (*P < 0.05, **P < 0.01, ***P < 0.001, ns= not significant using Welch’s t-test). Activity is lost after boiling and shown as a negative control for each sample. **c-d**, Expression (measured by qPCR) of *chiA* or *2345* in wild-type and mutant lines in mid-exponential growth in relation to the housekeeping gene, *rnpB*, in natural seawater-based Pro99 medium in presence and absence of colloidal chitin.

Once the recovered mutant lines showed growth rates similar to WT (Fig. S3), we tested the endochitinase and chitobiosidase activities reported previously in the WT^2^. We amended our samples with either colloidal chitin or chitosan, a form of chitin that is solubilized through partial deacetylation. Both additions were equally effective in stimulating the chitinase activity in WT cell-free spent media, and the activity disappeared upon boiling the samples (Fig. 2a,b) – consistent with the production of extracellular chitinase enzymes denatured upon heating. *ΔchiA* samples displayed neither endochitinase nor chitobiosidase activity, demonstrating that ChiA is required for both chitinase activity and true chitinase (Fig. 2a,b). The *ΔchiA* line showed higher expression of *2345* than in the WT (Fig.2d). Similarly, expression of *chiA* was higher in *Δ2345* than in the WT (Fig.2c), suggesting that cells lacking one gene compensate by expressing more of the other, which is often the case when proteins work in complexes or have similar functions^15^.

We note that despite the higher expression of the chitinase gene *chiA*, chitinase activity detected in *Δ2345* was significantly lower than in the WT (Fig. 2a,b). This result suggests that while not essential for the enzymatic activity, 2345 helps the ChiA enzyme perform the activity and that in its absence, the activity carries on less efficiently. Similarly, chitinase activity in the knock-down line *Δ2069* was reduced compared to the WT (Fig. S4). However, in this line, also expression of the chitinase enzyme *chiA* was reduced (Fig. S2c). Therefore, the reduced chitinase activity in *Δ2069* is due to a lower expression of c*hiA*, which also results in a higher expression of *2345*, like in *ΔchiA*. Due to *chiA* proximity to *2069* in the genome (Fig. S2a), the perturbation in *chiA* expression may be due to a disruption in the regulatory region of *chiA* that occurred while obtaining the edited line with the CRISPR-Cpf1 system. To test whether 2345 contributes to the chitinase activity by physically binding to ChiA forming complexes, we generated pETM-11-derived vectors to express these genes in *E*.*coli* in order to perform an *in vitro* analysis of the proteins. However, the expression of these genes is lethal to *E. coli*, so testing this hypothesis *in vitro* was not possible, as no viable colonies were obtained.

Putative chitinases containing chitin-binding domains but lacking glycosyl hydrolase domains have also been described in *Vibrio* and *Serratia* genera as possible adhesins or chitinase regulatory proteins^16,17^. Similarly to our results in *Synechococcus*, their production was induced by presence of chitin^16,18^, and no chitin degradation activity is attributed directly to them^19^. In *Vibrio*, deletion of CBP, a chitin-binding protein, results in a mutant expressing chitinolytic genes constitutively^20^. Likewise, the expression of *chiA* in *Synechococcus*, constitutively expressed in the WT, was found overexpressed in the *Δ2345* mutant line (Fig. 2c), suggesting that 2345 also regulates the expression of *chiA*.

Finally, because CBPs have been shown to facilitate chitin colonization in *V. cholerae*,^*21–23*^ we wondered whether *chiA* and *2345* had a similar role in *Synechococcus*. We tested this indirectly by estimating cell adhesion to added colloidal chitin – measuring both bulk fluorescence and cell number in suspension – in the WT and the loss of function mutants (Fig. S5). We used *Prochlorococcus* MED4 as a control, as it does not attach to chitin^2^. In all samples, the growth rate calculated based on the bulk fluorescence was not affected by the addition of colloidal chitin (Fig.S5a-d). But all *Synechococcus* lines (WT and mutants) amended with colloidal chitin showed a significant decrease in cell count in suspension by day 4 (Fig. S5 e-h). Cell loss in this planktonic state is due to cells attaching to the chitin polymer. Cells attached contribute to the fluorescence measured but cannot be detected by flow cytometry. We note that there was no appreciable difference in attachment between WT, *ΔchiA*, and *Δ2345* (Fig S5e-g), indicating that either the products of these genes are not involved in attachment or that chitin binding is multifactorial in *Synechococcus*. These results are consistent with previous findings showing that *Synechococcus* WH7803 can adhere to other surfaces^2^.

In summary, we show that the CRISPR-Cpf1 system can be delivered via electroporation in *Synechococcus marinus* to generate loss-of-function mutants. We identified *chiA* as the gene required for chitin degradation and 2345 as a chitinase-like protein indirectly involved in its activity. A major bottleneck in better understanding these minimal phototrophs’ physiology is the inability to easily manipulate the cells genetically. This work takes a significant step forward in obtaining a reliable toolbox for *Synechoccoccus* and, potentially, *Prochlorococcus*.

## Methods

### Culture conditions and growth curves

*Synechococcus* cells were grown under constant light flux at 12 μmol quanta m^−2^ s^−1^ and 24°C in natural seawater-based Pro99 medium containing 0.2-μm-filtered Sargasso Sea water, amended with Pro99 nutrients (N, P, and trace metals) prepared as previously described^24^. Where indicated, the samples were amended with high molecular weight chitosan or colloidal chitin (Millipore Sigma) to a final concentration of 56 μg/ml.

Growth was monitored using bulk culture fluorescence measured with a 10AU fluorometer (Turner Designs). Cell concentration was measured using a Guava easyCyte 12HT flow cytometer (EMD Millipore, Billerica, MA). Cells were excited with a blue 488 nm laser for measuring chlorophyll fluorescence (692/40 nm).

### Quantitative PCR analysis

*Synechococcus* cells grown at 12 μmol photons m^−2^ s^−1^ were collected by centrifugation. RNA samples were extracted with a standard acidic Phenol:Chloroform protocol and measured with Nanodrop (Thermo Scientific). RevertAid First Strand cDNA Synthesis Kit (Thermo Scientific) with random primers was used to obtain cDNA. Quantitative PCR reactions were performed in a CFX96 thermocycler (Bio-Rad) using the primers listed in Table S2. The expression of *rnpB* gene diluted 1:100 was used to normalize the results.

### Chitinase assay

*Synechococcus* WH7803 wild type and mutant cultures were grown in constant light at 12 μmol quanta m^−2^ s^−1^ in Pro99 media amended with high molecular weight chitosan or colloidal chitin (Millipore Sigma) to a final concentration of 56 μg/ml in triplicates. Cell concentration was measured using a Guava easyCyte 12HT flow cytometer (EMD Millipore, Billerica, MA). Cells were excited with a blue 488 nm laser for measuring chlorophyll fluorescence (692/40 nm). A volume containing 2E+09 total cell number was calculated and then centrifuged to remove cells from the spent media.

The supernatant was filtered through a 0.2 μm filter and concentrated using 30kDa Amicon® Ultra-15 Centrifugal Filter Units (Millipore) to a volume of 1,5 ml. Half the sample volume was boiled at 90°C for 30 minutes to serve as control. Each sample was then divided into 3 aliquots. Each aliquot was tested with one of the 3 substrates contained in the Chitinase kit (Sigma): 4-Methylumbelliferyl N,N′-diacetyl-β-D-chitobioside (substrate suitable for exochitinase activity detection or chitobiosidase activity), 4-Methylumbelliferyl N-acetyl-β-D-glucosaminide (substrate suitable for exochitinase activity detection of β-N-acetylglucosaminidase activity) and 4-Methylumbelliferyl β-D-N,N′,N′′-triacetylchitotriose (substrate suitable for endochitinase activity detection). The aliquots were mixed with these substrates and kept in darkness. The fluorescence of the 4-methyl-umbelliferone released by the chitinase activity in the sample was measured every 2 hours on a plate reader set at excitation 360 nm and emission at 450 nm.

### Electroporation and CRISPR plasmid construction

We constructed our vectors using the plasmid pSL2680 and designed the sgRNAs as described^10^. A homologous repair template was synthesized as left and right fragments with 700-750 bp of homology to each gene’s upstream and downstream sequences using the primers listed in Table S2. Cells in late-exponential phase (∼10^8 cell/ml) were pelleted and washed twice in ice-cold osmoprotectant (0.4M mannitol, 1mM HEPES pH 7.5) to remove all traces of seawater. Cells were then concentrated in 80μL (∼10^10 cells/ml) to which the plasmid of interest was added. Samples were electroporated (2.5 kv, 500 ohms and 25 μF) and resuspended in seawater media. After incubating for 24 hours at 10 μE m-2 s-1, cells were collected by centrifugation and pour-plated in sterile seawater based 0.3% low melting point agarose solution heated at 30°C with the addition of 50μM kanamycin sulfate, 10mM sodium bicarbonate, and 1mM sodium sulfite. Plates were transferred to ambient light conditions (12-15 μE m-2 s-1). Colonies were PCR screened for presence of appropriate deletions. Two rounds of plating or dilution to extinction were performed to obtain fully edited lines.

### In vitro expression vectors

*chiA, 2345* and *2069* full-length coding sequences were cloned into a pUC19-derived propagation vector for Golden Gate assembly. pETM-11 derived vectors for protein induction were then constructed to induce the expression of the three genes in BL21(DE3) or DH5alpha *E. coli* strains. However, cells could not grow when any of these proteins were induced.

## Supporting information

Plasmid maps

## Supplementary Figures

**Fig.S1:**
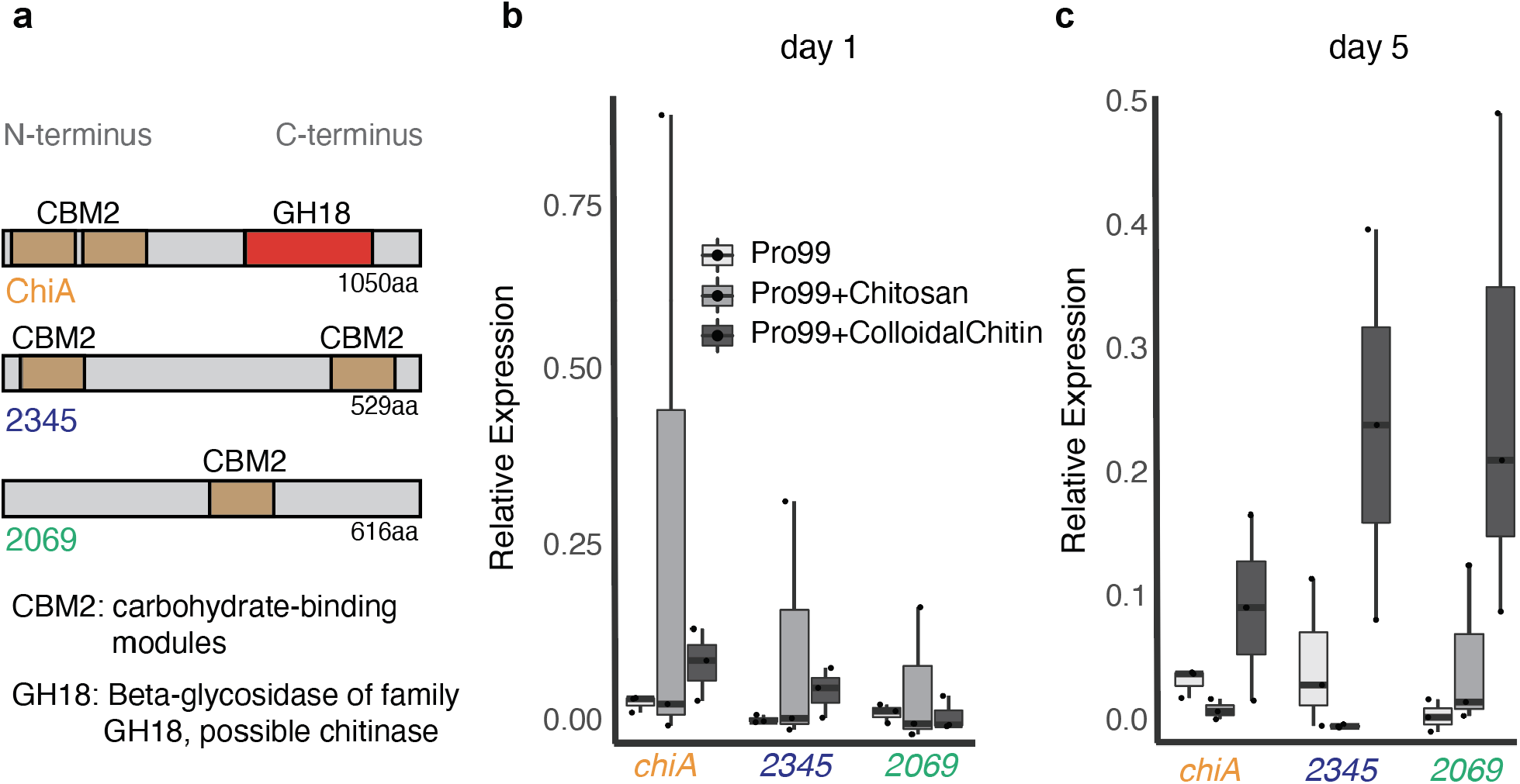
Identification of the genes of interest. **a**, Cartoon representation of domain structure of relevant exoproteins detected in marine *Synechococcus* WH7803. The number of amino acids is reported for each. Carbohydrate-binding modules (CMB2) are reported in brown, and Beta-glycosidase modules (GH18) in red. **b-c**, Expression (measured by qPCR) of *chiA, 2345* or *2069* in wild-type in mid-exponential growth in relation to the housekeeping gene, *rnpB*, in natural seawater-based Pro99 medium in presence and absence of colloidal chitin or chitosan.

**Fig.S2:**
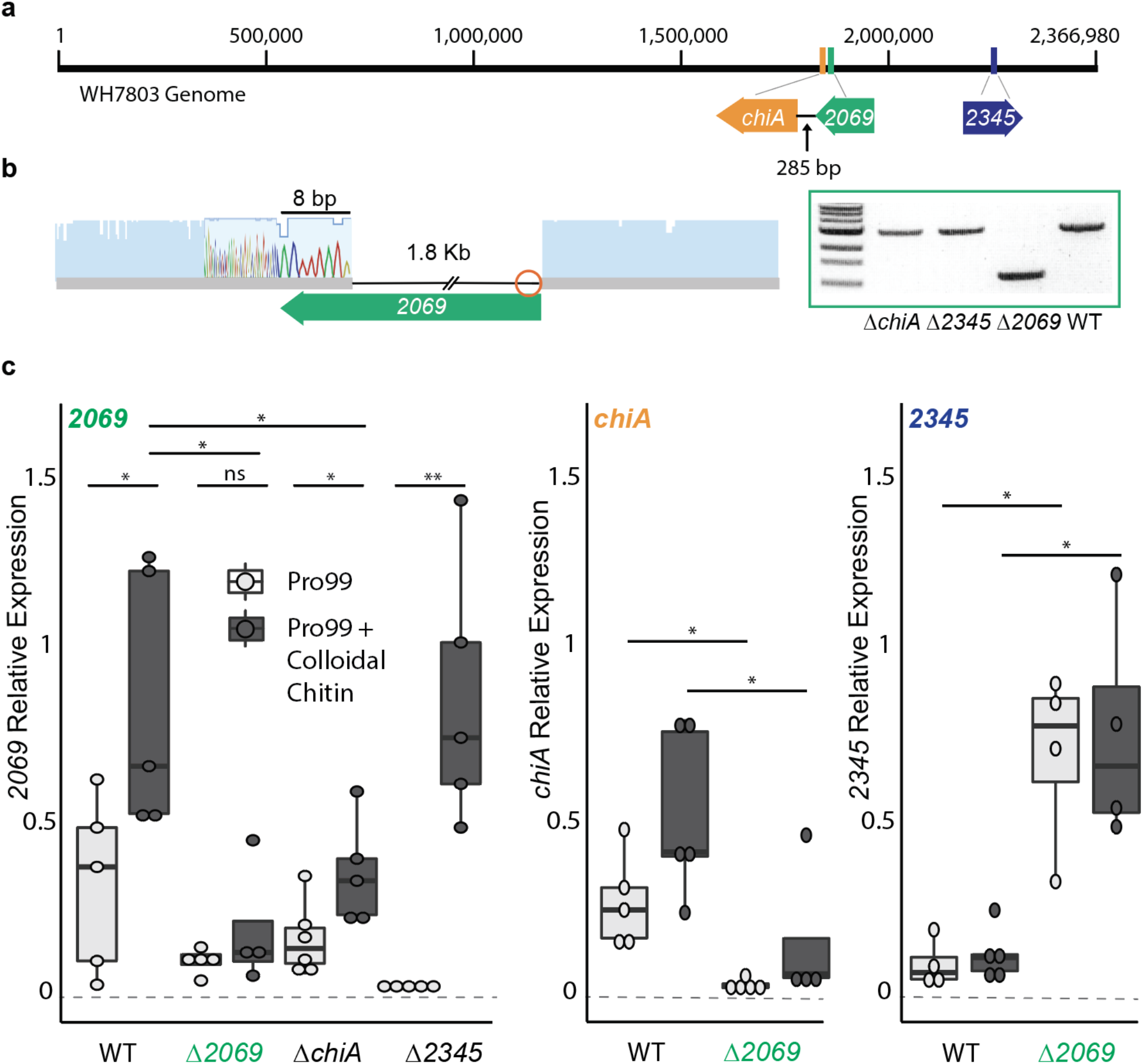
Mutant lacking *2069*. **a**, Cartoon representation of the WH7803 genome and the relative positions of all the three genes of interest. **b**, Schematic representation of the edited cell line obtained with the CRISPR-Cpf1 tool. Sanger sequences show the details of each deletion. Orange circles show the location of the PAM sites. PCR products indicate the length of each amplification using primers listed in Table S2. **c**, Expression (measured by qPCR) of *2069, chiA*, or *2345* in wild-type and mutant lines in mid-exponential growth in relation to the housekeeping gene, *rnpB*, in natural seawater-based Pro99 medium in presence and absence of colloidal chitin (*P < 0.05, **P < 0.01, ns= not significant using Welch’s t-test).

**Fig.S3:**
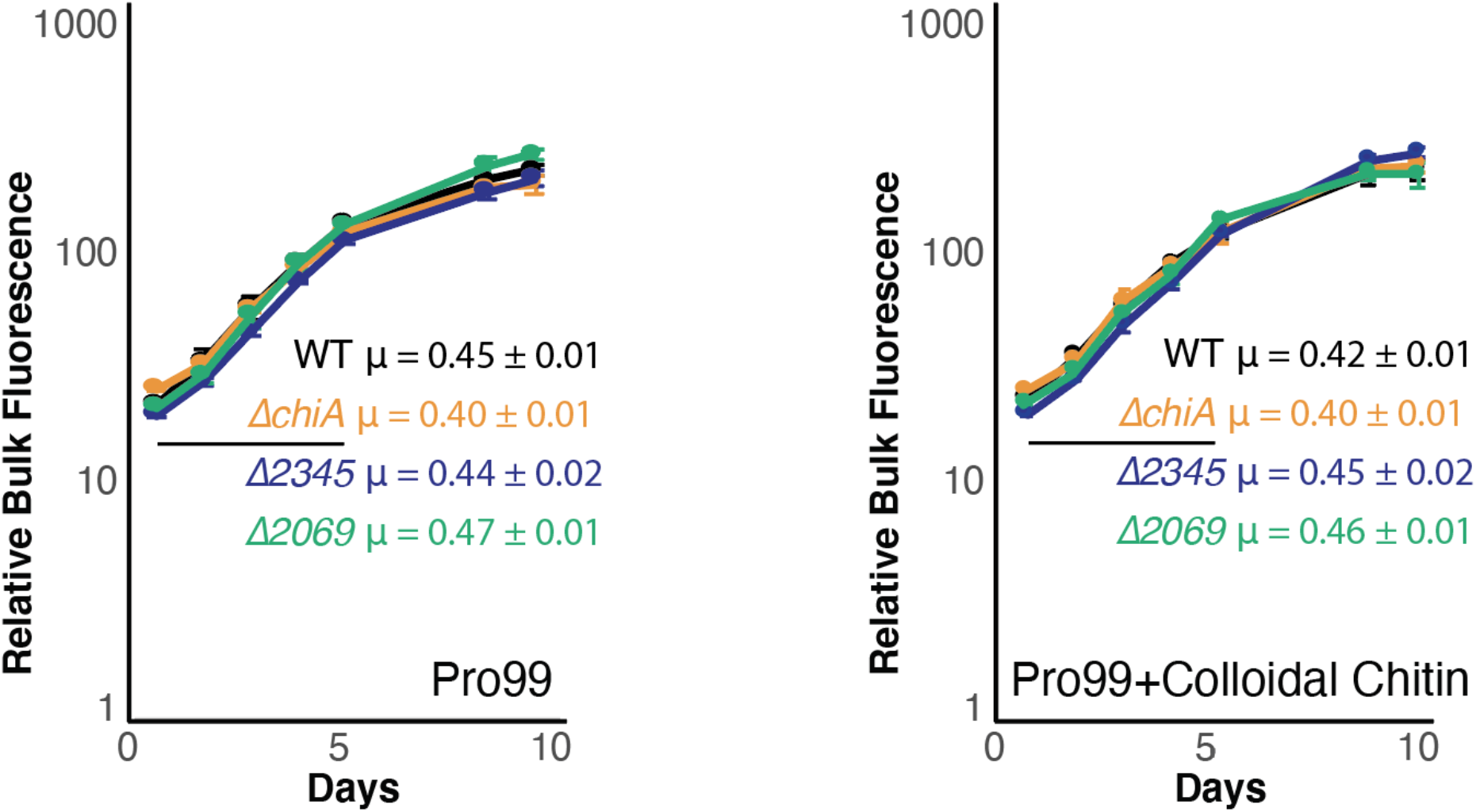
Growth rates in *Synechococcus* WT and mutant lines. Growth of *Synecococcus* WH7803 WT and recovered mutant lines in continuous light (at 12 μmol photons m^−2^ s^−1^) monitored by relative bulk culture chlorophyll fluorescence. Different colors show the average growth for each line in Pro99 media or Pro99 media amended with colloidal chitin. Growth rates and associated standard deviation (μ, in units day^-1^) was calculated in exponential phase (marked with a black line) and is shown for each curve.

**Fig.S4:**
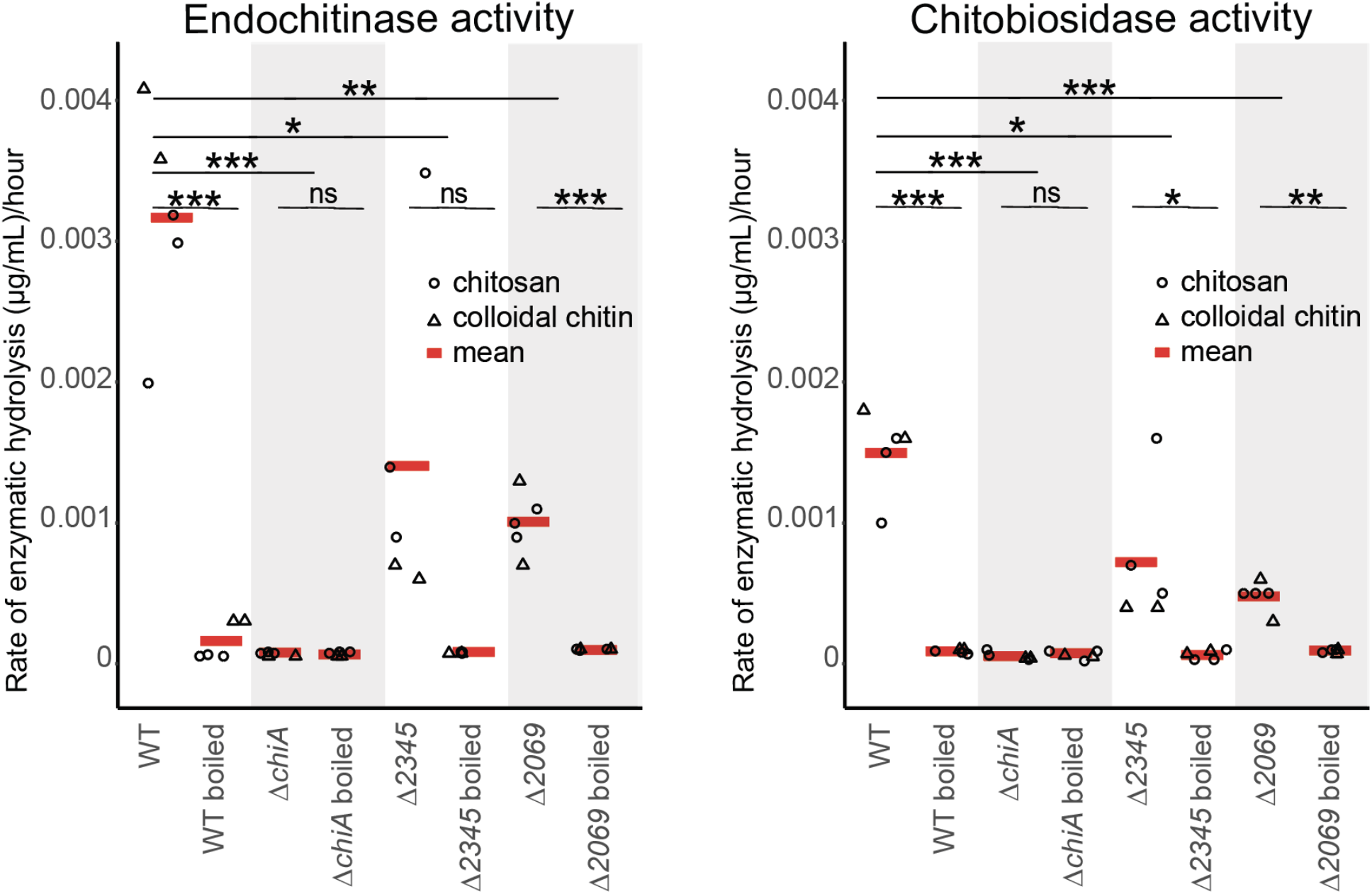
Chitin degradation activity. Endochitinase and Exochitinase (chitobiosidase) activities measured in wild-type and mutant lines spent media amended with either colloidal chitin or chitosan to a final concentration of 56 μg/ml (*P < 0.05, **P < 0.01, ***P < 0.001, ns= not significant using Welch’s t-test). Activity is lost after boiling and shown as a negative control for each sample. Data relative to WT, *ΔchiA*, and *Δ2345* is also reported in Figure 2a-b.

**Fig.S5:**
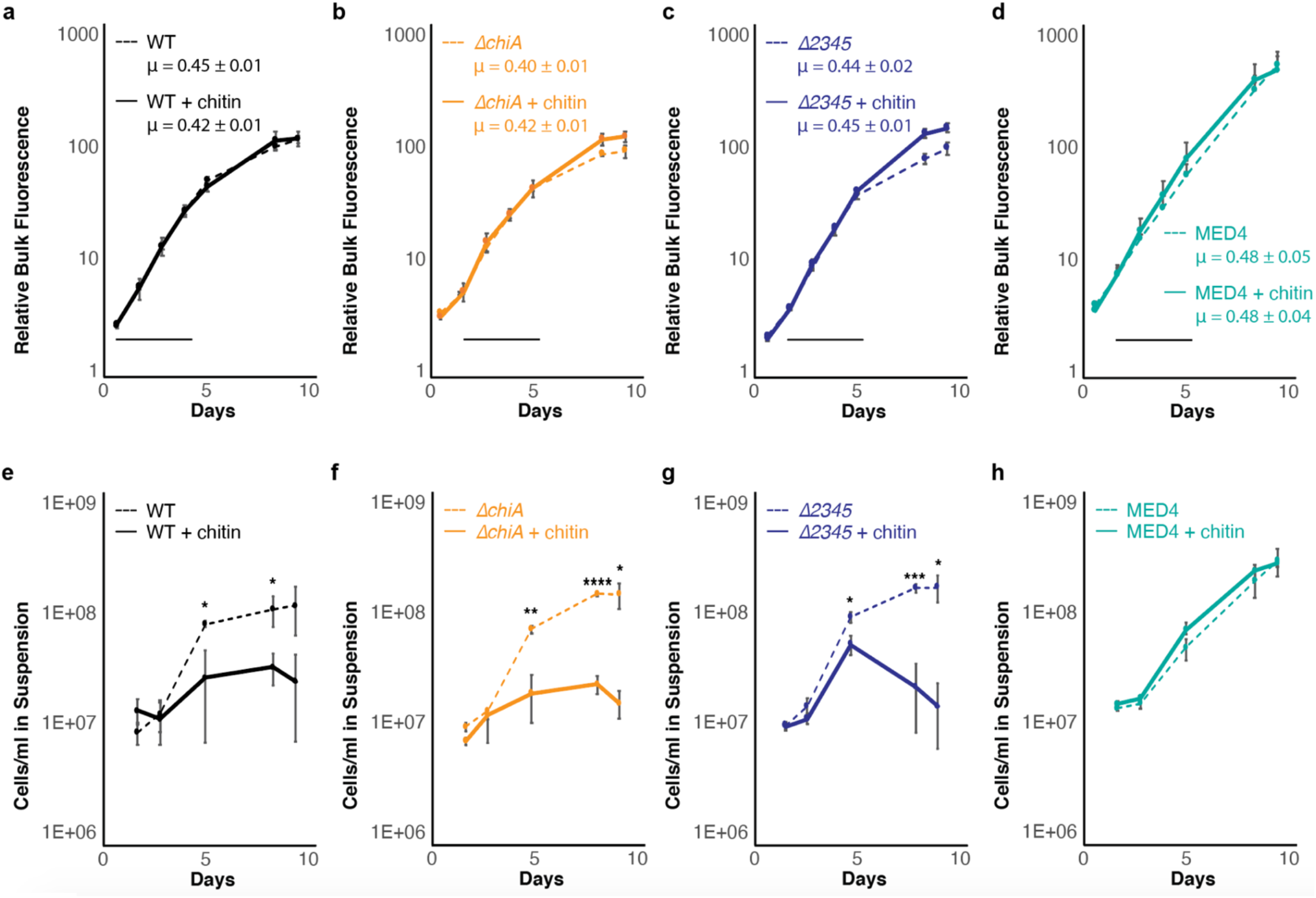
Adhesion of cells to colloidal chitin particles. Cultures were grown in Pro99 media with (solid line) and without (dashed line) added colloidal chitin. Growth was monitored by bulk chlorophyll fluorescence **a-d**, and cells in suspension were measured using flow cytometry **e-h**. Error bars show standard deviation between three biological replicates. The average growth rate and associated standard deviation (μ, in units day^-1^) was calculated in exponential phase (marked with a black line) and is shown for each curve. While growth rates between treatments did not differ, *Synechococcus* cells in suspension were less abundant in presence of chitin, as cells attaching to the polymer avoid detection via flow-cytometer. MED4, a high-light adapted *Prochlorococcus* ecotype, does not stick to chitin^2^, and consistently no difference was found in cells in suspension between the two treatments. (*P < 0.05, **P < 0.01, ***P < 0.001, ****P < 0.0001 using Welch’s t-test)

**Table S1: List of plasmids used**

pSL2680_*chiA* KanR CRISPR/Cpf1 plasmid containing gRNA and homologous repair template designed to target *chiA*

pSL2680_*2345* KanR CRISPR/Cpf1 plasmid containing gRNA and homologous repair template designed to target *2345*

pSL2680_*2069* KanR CRISPR/Cpf1 plasmid containing gRNA and homologous repair template designed to target *2069*

**Table S2:**
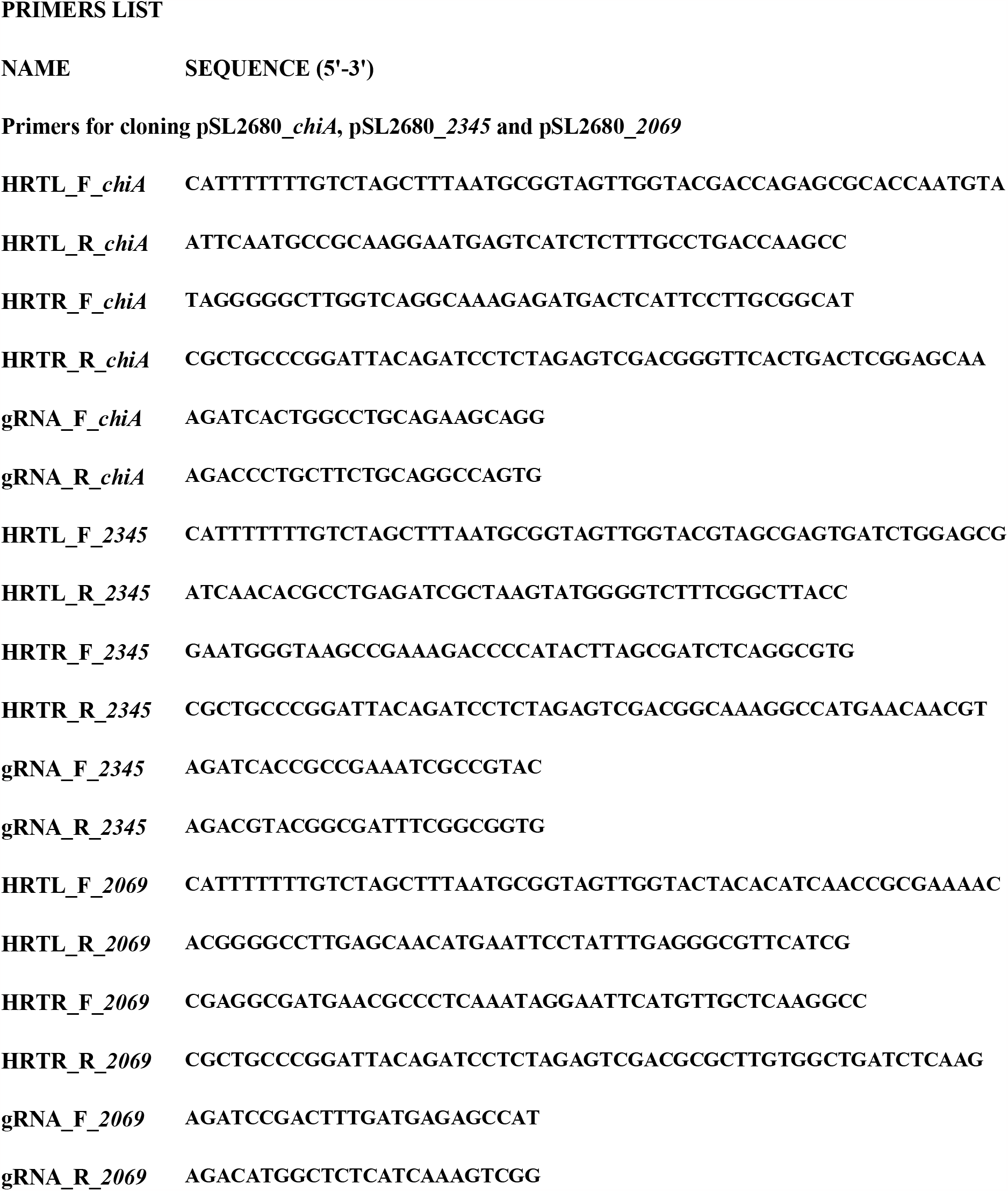
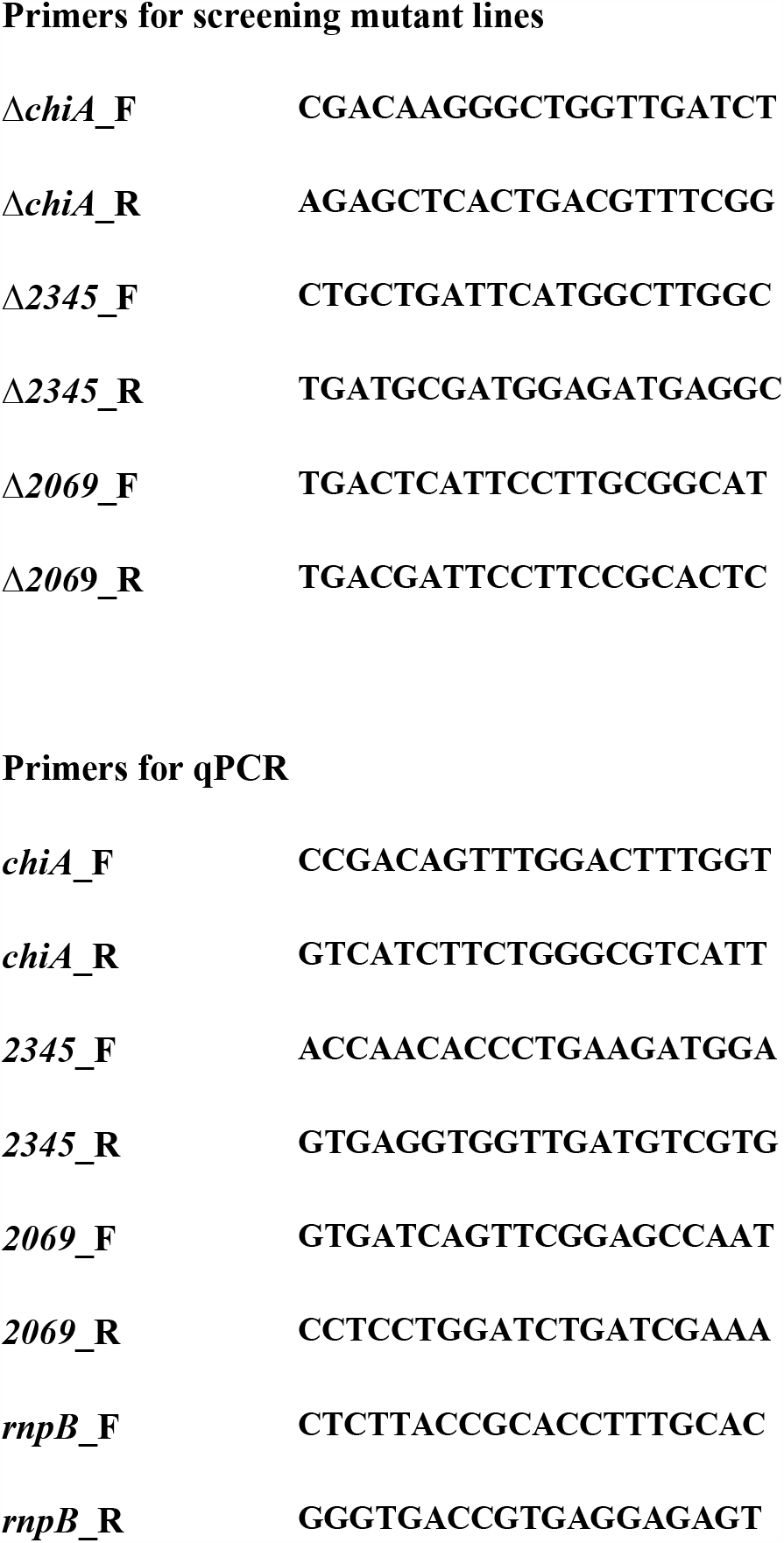
List of primers used.

## Acknowledgments

This research was supported in part by grants from the National Science Foundation (NSF-EDGE – 1645061 to SWC and NSF-EDGE 2035181 to SWC and Burton, B) and from the Simons Foundation (Life Sciences Award IDs 337262, 647135, 736564 to S.W.C.; SCOPE Award ID 329108 and 721246 to S.W.C.). G.C. was also supported by the Human Frontier Science Program (LT000069/2019-L). This paper is a contribution from the Simons Collaboration on Ocean Processes and Ecology (SCOPE).

## Contributions

G.C. designed experiments.

G.C., K.G.C., S.C., S.M.K. and D.M.K. performed experiments.

G.C and S.W.C. interpreted data.

G.C., and S.W.C. wrote the paper with contributions from all authors.

